# Tachykinin1-expressing neurons in the parasubthalamic nucleus control active avoidance learning

**DOI:** 10.1101/2024.11.03.621743

**Authors:** Ruining Hu, Nannan Wu, Tong Liu, Liuting Zou, Songjie Lv, Xiao Huang, Rongfeng K. Hu

**Author notes:** These authors contributed equally (Ruining Hu, Nannan Wu, Tong Liu).

## Abstract

Active avoidance is a type of instrumental behavior that requires an organism actively to engage in specific actions to avoid or escape from a potentially aversive stimulus and is crucial for the survival and well-being of organisms. It requires a widely distributed, hard-wired neural circuits spanning multiple brain regions, including the amygdala and thalamus. However, less is known about whether and how the hypothalamus encodes and controls active avoidance learning. Here we identify a previously unknown role for the parasubthalamic nucleus (PSTN), located in the lateral subdivision of the posterior hypothalamus, in the encoding and control of active avoidance learning. Fiber photometry calcium imaging shows that the activity of tachykinin1-expressing PSTN (PSTN^Tac1^) neurons progressively increases during this learning. Cell-type specific ablation and optogenetic inhibition of PSTN^Tac1^ neurons attenuates active avoidance learning, whereas optogenetic activation of these cells promotes this learning via a negative motivational drive. Moreover, the PSTN mediates this learning differentially through its downstream targets. Together, this study identifies the PSTN as a new member of the neural networks involved in active avoidance learning and offers us potential implications for therapeutic interventions targeting anxiety disorders and other conditions involving maladaptive avoidance learning.

## Introduction

Aversive learning is a type of learning that occurs when a neutral stimulus becomes associated with a negative or aversive outcome^1–5^ and plays a vital role for the survival of animals across various species^1–4,6^. It involves two fundamental types of conditioning: Pavlovian conditioning and instrumental conditioning^1–4^. The underlying neuronal circuits associated with aversive learning are complex and have been extensively studied in the context of fear conditioning, particularly through Pavlovian conditioning paradigms^7,8^. However, the neural mechanisms underlying instrumental defensive behaviors, such as active avoidance, are not completely understood. Indeed, several interconnected brain regions, such as the paraventricular thalamic nucleus (PVT)^9^, the lateral habenula (LHb)^10^, the amygdala^11–14^, the prefrontal cortex (PFC)^6,15^ and the periaqueductal gray (PAG)^16^, have been recently identified to involve active avoidance behavior, greatly expanding our understanding of the neural circuit mechanisms that underlie this behavior. In contrast, both traditional lesion and pharmacological experiments have found that the hypothalamus plays essential roles in regulating active avoidance^17–21^, but less is known about what hypothalamic neuronal types and their projections are involved in the regulation of this behavior and how these neurons and circuits are dynamically modulated by learning.

The parasubthalamic nucleus (PSTN), located in the lateral subdivision of the posterior hypothalamus^22,23^, is composed of mostly glutamatergic neurons, some of which express Tachykinin 1(*Tac1*), adenylate cyclase-activating polypeptide (*Adcyap1*) as well as corticotropin-releasing factor (*Crh*). Based on recordings and manipulations of these genetically labeled subpopulations of the PSTN, this brain region has recently been implicated in the regulation of feeding behaviors^24–27^, appetite suppression^28,29^, alcohol consumption^30–33^, arousal^34^, and fear-associated behaviors^26,35,36^. In addition, the PSTN has been found to promote conditioned taste aversion^22,37,38^ and place avoidance behaviors^26,28,35,36^. For instance, optogenetic activation of glutamatergic neurons or PACAP-expressing neurons or Tac1-expressing neurons in the PSTN promotes real-time place aversion^26,28,39,40^, suggesting that activation of the PSTN could generate a negative emotional state. Moreover, previous work has demonstrated that the PSTN sends outputs to the PVT^29,41^, an important brain node implicated in active avoidance learning ^9,42^, a process that can be prompted by negative motivational drives and be modified through experience ^3,4^. These results led us to hypothesize that the PSTN might be an attractive candidate brain node for the regulation of active avoidance learning. However, to date, no study has been directly examined the exact functional role of and *in vivo* neuronal responses of the PSTN as well as its associated circuits in active avoidance learning.

## Results

### Neural responses of PSTN^Tac^^1^ neurons during active avoidance learning

Based on recent studies supporting that *Tac1* gene represents the most useful genetic marker to label a subpopulation of PSTN glutamatergic neurons ^22,24,27,29,30,32,33,39^, we thus selected the Tac1-Cre transgenic mouse line for subsequent experiments. In this mouse line, Cre expression is under the control of the promoter for Tachykinin1(*Tac1*), allowing us to specially monitor and manipulate Tac1-labeled neurons’ activity. We first used fiber photometry, an approach that combines fiber optics and photometry to measure bulk neural dynamics in live animals^43^, to monitor the response of Tac1-expressing cells in the PSTN (PSTN^Tac1^) from male mice by direct foot shock. To achieve this goal, we unilaterally injected AAV expressing Cre-dependent fluorescent calcium indicator GCaMP7f in the PSTN of Tac1-IRES-Cre mice and in the same surgery installed a fiber optic above the viral injection site (Figure 1A). We observed a significant fluorescence increase in GCaMP7f-expressing mice rather than GFP-expressing controls while undergoing foot shock (Figure S1A and B). Post hoc analysis suggested that GCaMP7f expression was largely restricted within the PSTN (Figure 1B).

**Figure 1.**
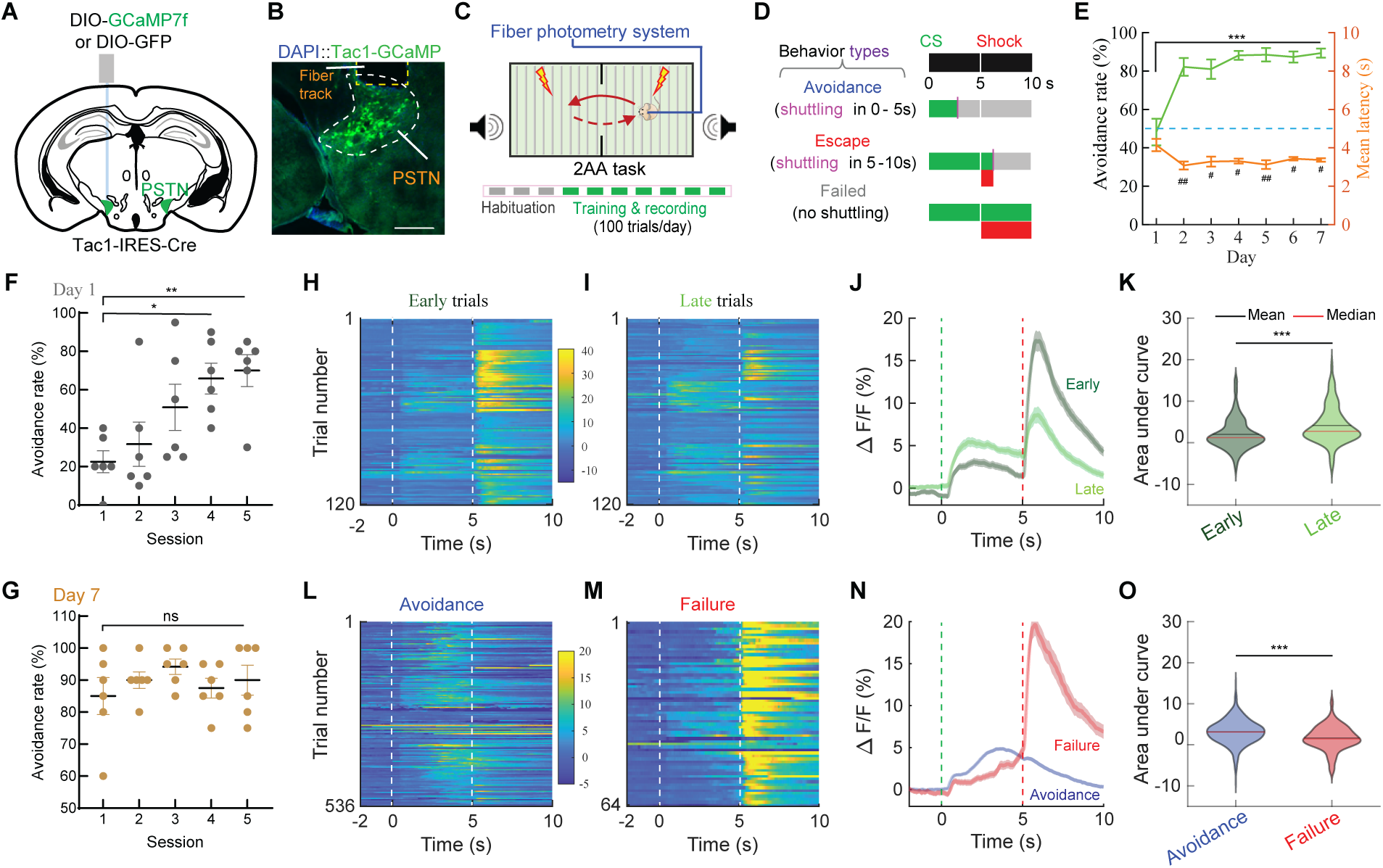
Neural dynamics of PSTN^Tac1^ neurons during active avoidance learning. (A) Schematic of Cre-dependent GCaMP7f injection into the PSTN of Tac1-IRES-Cre mice. (B) Example image showing GCaMP7f expression and fiber location in the PSTN. Yellow dashed line shows fiber track; white dashed line delineates the PSTN. Scale bar, 200 µm. (C) Schematic of two-chamber active avoidance (2AA) task (upper) and the training pipeline (bottom). Red arrow lines refer to shuttle direction; Gray boxes and green boxes refer to 3-day habituation and 7-day training and recording, respectively. (D) Behavioral protocol and definition of different behavioral readouts. In 2AA, a test mouse experienced a tone (Conditioned stimulus, CS) followed by an aversive foot shock (Unconditioned stimulus, US). Each trial has a constant 10-second-long time window and the trial-trial interval is ∼ 15–30 s. According to the shuttle onset in a given trial, it can be divided into three types: an avoidance trial (shuttle within 0–5 s and no shock), an escape trial (shuttle within 5–10 s and a short foot shock) and a failed trial (no shuttle and a maximum of shock). Green-filled boxes, CS duration; red-filled boxes, shock duration; purple lines, shuttle onset. (E) Time course and line graph illustrating average avoidance rate and mean latency along 7-day training procedure. One-way repeated-measures ANOVA with Bonferroni post-hoc correction (For avoidance rate: ***P < 0.001; For mean latency: ^#^P < 0.05, ^##^P < 0.01). (F, G) Scatterplots illustrating average avoidance rate across five sessions (20 trials/session) on day 1 (F) and on day 7 (G). One-way repeated-measures ANOVA with Bonferroni post-hoc correction (*P < 0.05 and **P < 0.01). (H, I) Heatmaps for pooling normalized fluorescence signals from the first 20 trials (early trials, H) and the last 20 trials (late trials, I) of each mouse. Time window for each trial is from 2 s before CS to 10 s after CS. Scale bar refers to % ΔF/F. (J) The peri-stimulus time histogram (PSTH) plots for all early trials and late trials during the time window. (K) Violin plots for the area under curve (AUC) of the average responses of early trials and late trials during CS. Two-tailed paired t test (***P < 0.001). (L, M) Heatmaps for pooling normalized fluorescence signals from avoidance trials (L) and failure trials (M) of each mouse. Time window for each trial is from 2 s before CS to 10 s after CS. Scale bar refers to % ΔF/F. (N) The PSTH plots for all avoidance trials and failure trials during the time window. O, Violin plots for the AUC of the average responses of avoidance trials and failure trials during CS. Two-tailed unpaired t test (***P < 0.001). In (E-O), n = 6 mice; in (E-K), n = 20 early trials and 20 late trials each animal; in (G), n = 5 sessions from 100 trials each animal; and in (L-O), n = 536 avoidance trials and n = 64 failure trials. In (E, F, G, J, K, N and O), data was shown as mean ± SEM; in (K, O), shown as median. For detailed statistics information, see Supplementary Table 1.

Next, to examine the *in vivo* neuronal responses of the PSTN in active avoidance learning, we used a modified two-way active avoidance learning task (2AA; Figure 1C and D), based on previous work^9^. In our study, the 2AA task included a 7-day training procedure (100 trials/day) (Figure 1C). During this experiment, a test mouse experienced a tone (Conditioned stimulus, CS), and must shuttle to the other side of chamber in a given response window to prevent an aversive foot shock (Unconditioned stimulus, US) (Figure 1D). The performance of active avoidance learning was calculated by the percentage of cue-guided successful shuttles over total trials (Avoidance rate) and the duration from trial onset to crossing compartments (Latency). Over the course of training, mice significantly increased their successful shuttling rates to avoid foot shock occurrence and decreased the latency to cross compartments (Figure 1E). Remarkably, the avoidance rate could reach and stay up to 70% across day 2 - day 7 (Figures 1E and S1H), suggesting that our modified 2AA training system is highly effective to being applied to investigate active avoidance learning. To further examine the avoidance performance on day 1, we sequentially and averagely divided 100-trial training into five sessions. We found that animals progressively improve their shuttling performance spinning from 22.5±5.7% to 70.0±8.3% (Figure 1F). Moreover, on day 7, mice could exhibit a stable behavioral performance (>70%) across sessions (Figure 1G).

Based on animals’ behavioral performance (Figure 1E-G), we then focused on analyzing calcium fluorescent transients on days 1 and 7. On day 1, we observed a significant increase in CS-evoked calcium fluorescent signals in late trials relative to those in early trials (Figure 1H-K), strongly indicating that active avoidance learning is capable of reshaping the activity of these cells. Next, to examine the neuronal dynamics for both avoidance trials and failure trials from well-trained mice on day 7, we pooled all those calcium imaging data and analyzed the response patterns for avoidance trials and failure trials, respectively. We observed a significant increase in CS-evoked calcium fluorescent signals in avoidance trials compared to those in failure trials (Figure 1L-O), indicating that well-learned mice apply a higher CS-evoked response in guiding avoidance behavior. In contrast, although our control mice also exhibited a considerable behavioral performance in 2AA task (Figure S1C), they had no bulk changes in fluorescence throughout the training (Figure 1D-G). Moreover, consistent with the behavioral performance during 2AA (Figures 1E and S1H, I), a very similar neuronal response pattern for those GCaMP-expressing male mice was observed on days 2-7 (Figure S1H-J). Finally, we also examined the response patterns of PSTN^Tac1^ neurons of female mice during 2AA. We analyzed behavioral performance and neuronal dynamics in female GCaMP-expressing mice on day 1, and found similar neuronal response patterns between early trials and late trials, compared to those in male mice, regardless of the behavioral performance (Figure S2). These data illustrate that the activity of PSTN^Tac1^ neurons in both sexes of mice can be shaped in a similar manner on the early stage of 2AA task.

Together, our study modified a new 2AA training system that can effectively and quickly evaluate the animals’ avoidance performance. Using this system, we provided the first *in vivo* calcium response evidence for modulation of PSTN^Tac1^ neurons’ activity by learning in both sexes.

### PSTN^Tac^^1^ neurons are required for mediating active avoidance learning

We next asked whether the activity of these cells is necessary to control active avoidance learning. To this goal, we conducted cell-specific ablation of PSTN^Tac1^ neurons by bilaterally injecting AAVs expressing Cre-dependent caspase-3 (or EGFP as a control) into the PSTN of Tac1-IRES-Cre mice (Figure 2A). Post hoc histological analysis demonstrated that ablation remarkably reduced the number of PSTN^Tac1^ neurons relative to that of control mice (Figure 2B-C). The majority of control mice displayed robust avoidance behavior in 2AA task with high avoidance rates and low latencies to trial onset (Figure 2D-I), but all ablated mice showed significantly attenuated active avoidance behaviors throughout the training (Figure 2F-I), suggesting that PSTN^Tac1^ neurons are required for active avoidance learning. We then tested whether PSTN^Tac1^ neurons’ activity is indispensable for the maintenance of learned active avoidance behavior (Figure S3). We screened a cohort of Tac1-IRES-Cre mice displaying robust avoidance performance (with an average avoidance rate > 70%) after 5-day 2AA training. These selected mice were then bilaterally injected with Cre-dependent caspase-3 (or EGFP as a control) into the PSTN. We found that ablated mice exhibited no differences in behavioral performance compared to those in control mice on post-test day 1 (Figure S3E, F). On post-test day 2, ablated mice and control mice had no significant difference in avoidance rate but had a small difference in the latencies to trial onset (Figure S3E, F). The results illustrated that ablation of PSTN^Tac1^ neurons largely had no effect on the learned active avoidance behavior. Collectively, these findings suggest that PSTN^Tac1^ neurons are required for active avoidance during the learning stage rather than post-learning stage.

**Figure 2.**
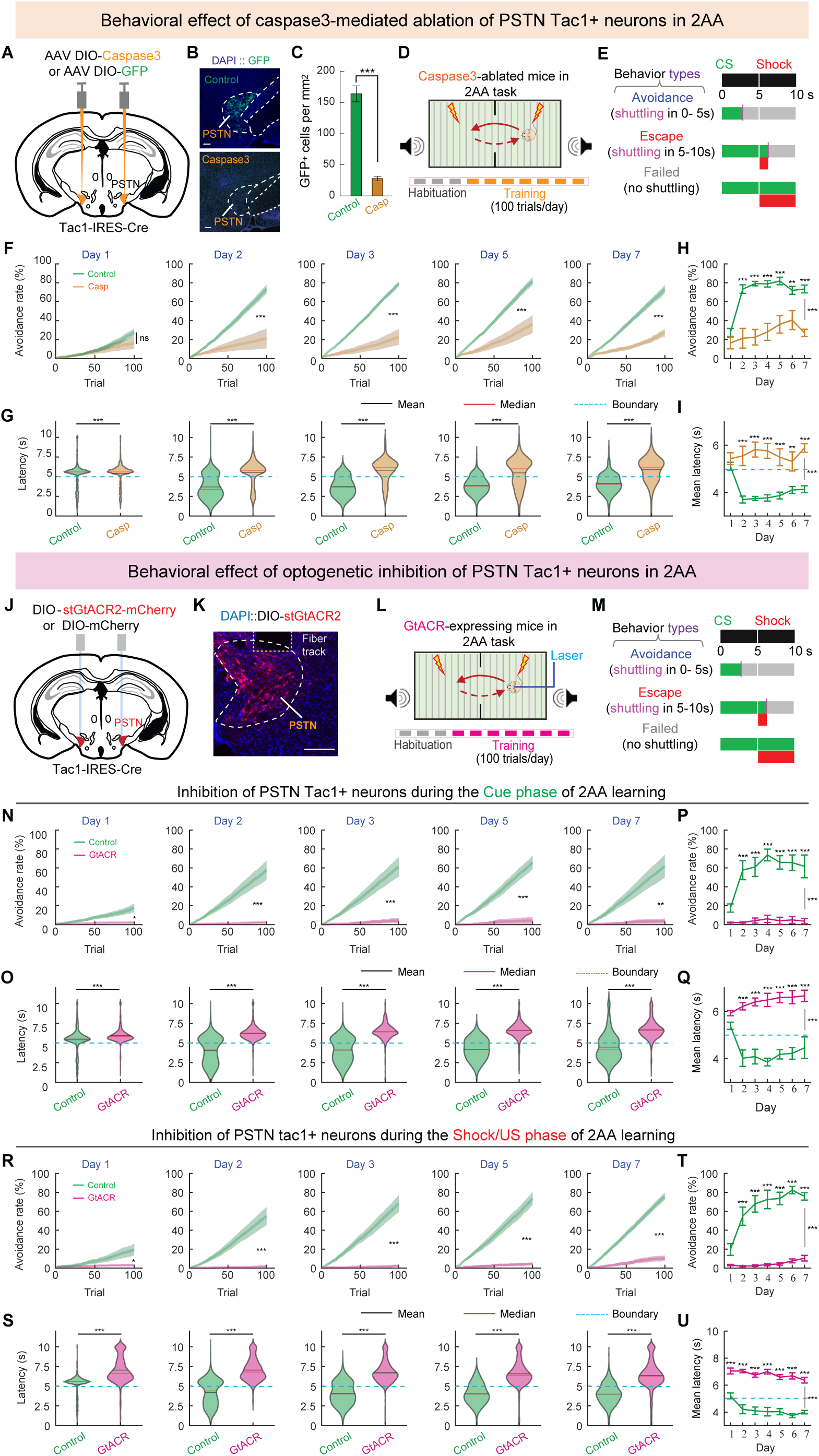
Necessity of PSTN^Tac1^ neurons mediating active avoidance learning. (A) Schematic of Cre-dependent caspase3 injection into the PSTN of Tac1-IRES-Cre mice. (B) Example images showing caspase injection (bottom) effectively ablated PSTN Tac1^+^ neurons whereas control GFP injection (upper) did not. Scale bar, 200 µm. (C) Bar graph illustrating caspase ablation significantly reduced the number of GFP^+^ cells in the PSTN compared to the controls. Two-tailed Mann Whitney test (***P < 0.001). (D) Schematic of two-chamber active avoidance task and training protocol. Gray boxes and orange boxes refer to 3-day habituation and 7-day training, respectively. (E) Behavioral protocol and definition of different behavioral readouts. (F, G) Average cumulative distribution of avoidance rates (F) and violin plots of latencies (G) for Caspase-ablated mice and EGFP-expressing controls on days 1-3, 5 and 7 of 2AA task. Two-way repeated-measures ANOVA with Bonferroni post-hoc correction in (F) (ns = not significant; ***P < 0.001) and two-tailed unpaired t test in (G) (***P < 0.001). (H, I) Line graphs of average avoidance rate (H) and mean latency (I) for caspase3-experessing mice and GFP-expressing controls along 7-day training. Two-way repeated-measures ANOVA with Bonferroni post-hoc correction (**P < 0.01 and ***P < 0.001). (J) Schematic showing viral injection and fiber implantation for optogenetic inhibition of PSTN Tac1^+^ neurons. (K) Example image of GtACR expression and fiber location in the PSTN. Yellow dashed line shows fiber track; white dashed line delineates the PSTN. Scale bar, 200 μm. (L) Schematic of two-chamber active avoidance task and training protocol. Gray boxes and purple boxes refer to 3-day habituation and 7-day training and inhibition, respectively. (M) Behavioral protocol and definition of different behavioral readouts. (N, O) Average cumulative distribution of avoidance rates (N) and violin plots of latencies (O) for GtACR-expressing mice and mCherry-expressing controls during CS inhibition on days 1-3, 5 and 7. Two-way repeated-measures ANOVA with Bonferroni post-hoc correction in (N) (*P < 0.05, **P < 0.01 and ***P < 0.001) and two-tailed unpaired t test in (O) (***P < 0.001). (P, Q) Line graphs of average avoidance rate (P) and mean latency (Q) for GtACR-expressing mice and mCherry-expressing controls along 7-day training. Two-way repeated-measures ANOVA with Bonferroni post-hoc correction (***P < 0.001). (R, S) Average cumulative distribution of avoidance rates (R) and violin plots of latencies (S) for GtACR-expressing mice and mCherry-expressing controls during US inhibition on days 1-3, 5 and 7. Two-way repeated-measures ANOVA with Bonferroni post-hoc correction in (R) (*P < 0.05 and ***P < 0.001) and two-tailed unpaired t test in (s) (***P < 0.001). (T, U) Line graphs of average avoidance rate (T) and latency (U) for GtACR-expressing mice and mCherry-expressing controls along 7-day training. Two-way repeated-measures ANOVA with Bonferroni post-hoc correction (***P < 0.001). In (F-I), n = 8 control mice and 8 Caspase-3 ablated mice; in (N-Q), n = 9 control mice and 7 GtACR-expressing mice; in (R-U), n = 8 control mice and 8 GtACR-expressing mice. In (F-I, N-Q, and R-U), data was shown as mean ± SEM; in (G, O and S), shown as median. For detailed statistics information, see Supplementary Table 1.

To further investigate the exact role of activities of PSTN^Tac1^ neurons during CS or US in controlling active avoidance behavior, we optogenetically suppressed the activity of these neurons bilaterally using a Cre-dependent silencing opsin GtACR2 (Figure 2J-M). We first examined the necessity of activity of these neurons during CS by delivering 5 s-long continuous photoinhibition during that phage while mice were performing in the 2AA task. Our results demonstrated that photoinhibition significantly attenuated active avoidance behavior in GtACR2-expressing mice, but not mCherry-expressing controls (Figure 2N-Q). Similarly, we then performed the experiment for optogenetic inhibition of PSTN^Tac1^ neurons during US. The results showed a similar behavioral effect with that in photoinhibition during CS (Figure 2R-U). Together, both ablation and photoinhibition experiments revealed that PSTN^Tac1^ neurons permit active avoidance learning.

### Optogenetic activation of PSTN^Tac^^1^ neurons promotes active avoidance learning and encodes aversive memory

Because our calcium imaging results found that the activity of PSTN^Tac1^ neurons progressively increases during 2AA learning (Figure 1H-K), we thus hypothesized that activation of these neurons during CS is sufficient to promote active avoidance behavior. To this end, we bilaterally injected a Cre-dependent activating opsin ChR2 into the PSTN of male Tac1-IRES-Cre mice (Figure 3A, B). 5 s pulsed light stimulation for activation of PSTN^Tac1^ neurons was delivered once a trial starts in the 2AA task (Figure 3C, D). Our results showed that optogenetic activation of these cells during CS significantly shortened the shuttle latencies and increased avoidance rates (Figure 3E-H), suggesting that activation of PSTN^Tac1^ neurons in male mice can promote active avoidance learning. Furthermore, we also examined behavioral effect by activation of these neurons in female mice, and found that optogenetic activation of PSTN^Tac1^ neurons in female mice significantly reduced the mean latencies to cross the other side (Figure S4) and showed an increased trend in their avoidance rates (Figure S4). This difference in avoidance rates between sexes may lie in greater individual variations in behavioral performance in females. Altogether, despite some sex differences in behavioral performance, optogenetic activation of PSTN^Tac1^ neurons in both sexes of mice has a promoting effect on animal’s active avoidance behavior.

**Figure 3.**
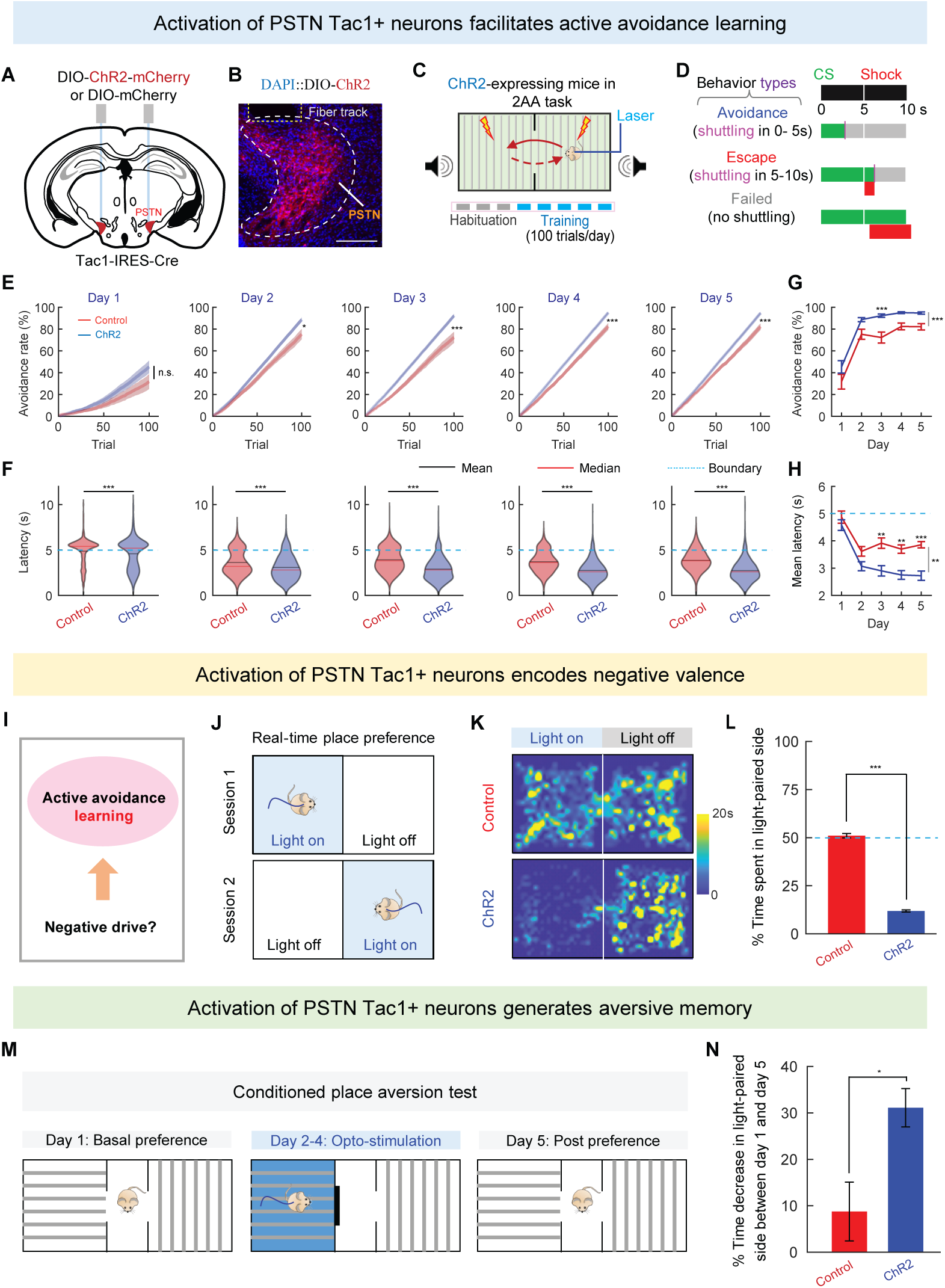
Sufficiency of PSTN^Tac1^ neurons mediating active avoidance learning. (A) Schematic showing viral injection and fiber implantation for optogenetic activation of PSTN Tac1^+^ neurons. (B) Example image of injection site and ChR2 expression in the PSTN of Tac1-IRES-Cre animals. Yellow dashed line shows fiber track; white dashed line delineates the PSTN. Scale bar, 200 μm. (C) Schematic of two-chamber active avoidance task and training pipeline. Gray boxes and blue boxes refer to 3-day habituation and 5-day training, respectively. (D) Behavioral protocol and definition of different behavioral readouts. (E, F) Average cumulative distribution of avoidance rates (E) and violin plots of latencies (F) for ChR2-experessing mice and mCherry-expressing controls on each day. Two-way repeated-measures with Bonferroni post-hoc correction in (E) (ns = not significant; *P < 0.05 and ***P < 0.001) and two-tailed unpaired t test in (F) (***P < 0.001). (G, H) Line graphs of average avoidance rate (G) and mean latency (H) for ChR2-experessing mice and mCherry-expressing controls along 5-day training. Two-way repeated-measures ANOVA with Bonferroni post-hoc correction (**P < 0.01 and ***P < 0.001). (I) Schematic of experiment model. (J) Schematic showing real-time place preference assay (RTPP). Light blue area indicates the chamber paired with light stimulation when the animal enters. (K) Representative heatmaps showing locomotion trajectories of control (top) and ChR2-experessing (bottom) mice in the RTPP test. (L) ChR2-expressing animals displayed a significant decrease in time spent in stimulation-coupled chamber compared to mCherry-expressing controls. Two-tailed Mann-Whitney test (***P < 0.001). (M) Schematic of conditioned place aversion (CPA) test and experimental procedures. (N) During post preference test on day 5, ChR2-expressing animals exhibited a remarkable reduction in time spent in preferred side on day 1 compared to mCherry-expressing controls, suggesting that activation of MeA GABAergic neurons could induce a long-term aversive memory for stimulation-coupled chamber. Two-tailed Mann-Whitney test (*P < 0.05). In (E-H), n = 8 control mice and 14 ChR2-expressing mice; in (L), n = 9 control mice and 9 ChR2-expressing mice; in (N), n = 11 control mice and 8 ChR2-expressing mice. In (E-H, L, and N), data was shown as mean ± SEM; in (F), shown as median. For detailed statistics information, see Supplementary Table 1.

As known, learning is highly dependent on individuals’ motivational drive^1,2,12,44,45^. Consistent with a recent study^40^, we used a real-time place preference assay and confirmed that animals expressing ChR2 into PSTN^Tac1^ neurons developed a consistent avoidance for the chamber coupled with light stimulation compared to the controls (Figure 3I-L), suggesting that PSTN^Tac1^ neurons encode negative motivational drive in promoting this behavior. To exclude the possibility of generating this aversive effect by sparse cells of PSTN neighboring areas, we conducted additional characterizations of behavioral effects of activation of sparsely Tac1-expressing neurons in two adjacent areas of PSTN, demonstrating no differences between light-coupled sides and its counter-balanced sides (Figure S5). These results indicate that PSTN^Tac1^ neurons, but not its adjacent Tac1-expressing neurons, encode negative motivational drive in promoting this behavior.

To further test whether optogenetic activation of PSTN^Tac1^ neurons could generate aversive memory, we applied a five-day conditioned place aversion assay, in which includes pre-testing baseline phase (day 1), stimulation phase (days 2-4) and post-testing phase (day 5) (Figure 3M). We found that optogenetic activation of these neurons from ChR2-expressing mice could significantly reduce the percentage of time spent on the stimulation-paired side between pre-testing baseline phase and post-testing phase (Figure 3N), which is divergent from a recent study^40^.

### Downstream targets of the PSTN mediate active avoidance learning

We next sought to dissect what the downstream circuitry of the PSTN is involved in mediating active avoidance learning. Indeed, a recent study has mapped and functionally characterized the whole-brain downstream targets of cell-type specific neurons in the PSTN in feeding behavior^29^ and the identified projections are largely consistent with those by traditional anterograde tracer experiment^23^. Here we used a viral-based anterograde tracing approach to confirm major projections from the PSTN (Figure S6), including the PVT/intermediodorsal nuclei of the dorsal thalamus (IMD) complex, the central nucleus of the amygdala (CeA), and the parabrachial nucleus (PBN). Among its projection targets, the PVT has been recently found to involve associative learning and defensive behaviors^9,42^. Therefore, we speculated that the PSTN-to-PVT/IMD pathway is an attractive candidate for mediation of active avoidance learning. To fully characterize the circuit dynamics and functional role of this circuit, we used axon-localized GCaMP6s based fiber photometry and optogenetics-assisted circuit mapping and manipulations (Figure 4A-D). Similar to the findings from the soma, the PSTN-to-PVT/IMD circuit signals aversive information on active avoidance behavior and is modulated by learning (Figure 4E-H). Moreover, optogenetic manipulations confirmed that this circuit is both necessary and sufficient to mediate this learning (Figure 4I-P). Together, the PVT/IMD is an important downstream target of the PSTN in the regulation of active avoidance learning, adding the PSTN as a new member to the established neural network that underlies this behavior.

**Figure 4.**
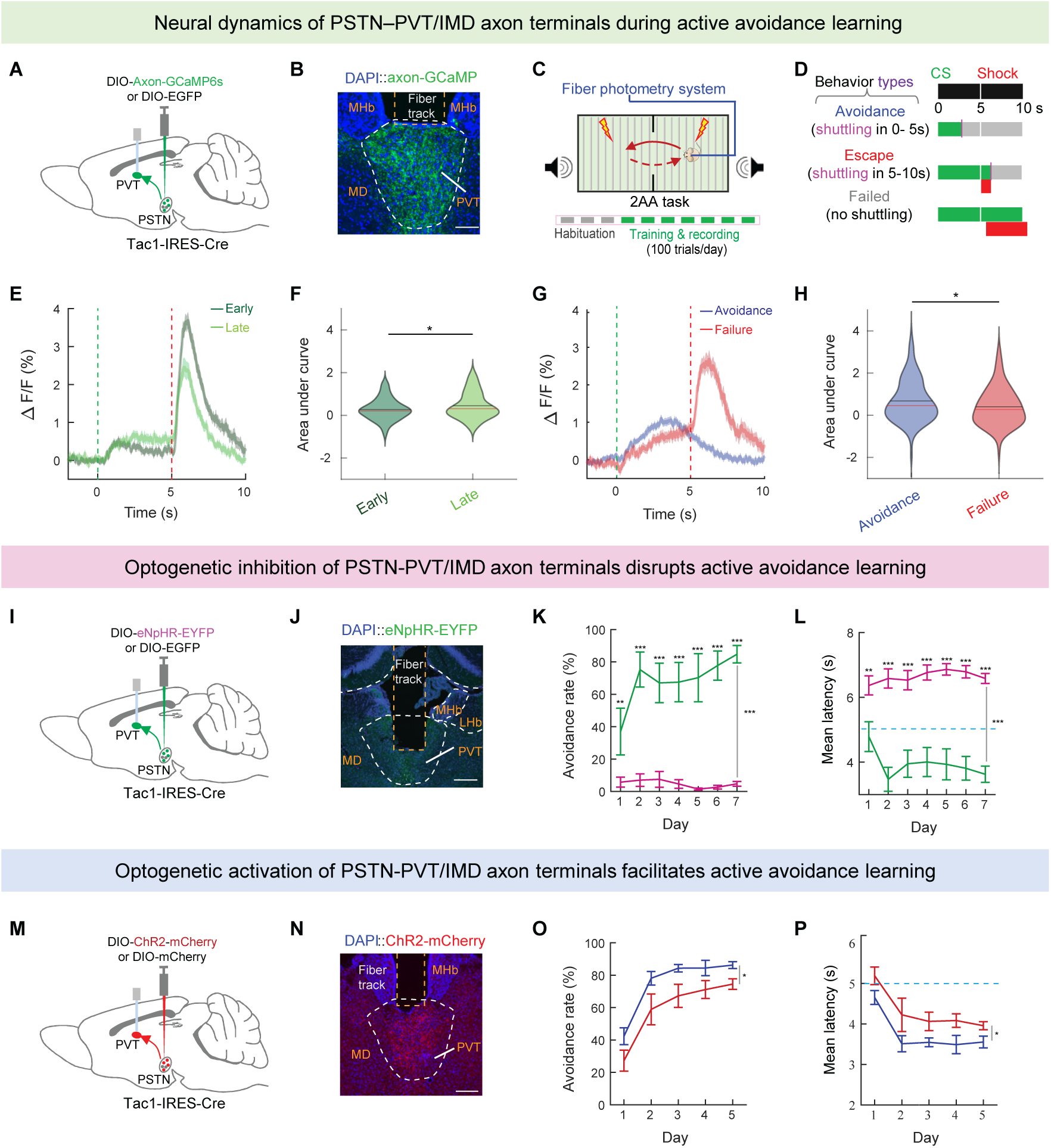
The PSTN projects to the PVT/IMD to mediate active avoidance learning. (A) Schematic of Cre-dependent axon GCaMP6s injection into the PSTN and fiber placement in the PVT. (B) Example image showing axon-GCaMP6s expression and fiber location in the PVT. Orange dashed line indicates fiber location and red dashed line delineates the PVT. Axon-GCaMP signals were strengthened by immunostaining with anti-GFP. Scale bar, 100 µm. (C) Schematic of two-chamber active avoidance (2AA) task (upper) and the training pipeline (bottom). (D) Behavioral protocol and definition of different behavioral readouts. (E) PSTH plots for all early trials and late trials during the time window. Time window for each trial is from 2 s before CS to 10 s after CS. (F) Violin plots for AUC of average responses of early trials and late trials during CS. Two-tailed paired t test (*P < 0.001). (G) PSTH plots for all avoidance trials and failure trials during the time window. Time window for each trial is from 2 s before CS to 10 s after CS. (H) Violin plots for AUC of average responses of avoidance trials and failure trials during CS. Two-tailed unpaired t test (*P < 0.001). (I) Schematic showing eNpHR injection and fiber implantation for optogenetic inhibition of PSTN-PVT circuit. (J) Example image of eNpHR expression in axon terminals in the PVT. Orange dashed line indicates fiber location and white dashed line delineates the PVT. Scale bar, 200 μm. (K, L) Line graphs of average avoidance rate (K) and latency (L) for eNpHR-expressing mice and mCherry-expressing controls along 7-day training. Two-way repeated-measures ANOVA with Bonferroni post-hoc correction (**P<0.01 and ***P < 0.001). (M) Schematic showing viral injection and fiber implantation for optogenetic activation of PSTN-PVT circuit. (N) Example image of axon terminal expression of ChR2 in the PVT. Orange dashed line indicates fiber location and red dashed line delineates the PVT. axon terminal mCherry signals were strengthened by immunostaining with anti-mCherry. Scale bar, 200 μm. (O, P) Line graphs of average avoidance rate (O) and mean latency (P) for ChR2-experessing mice and mCherry-expressing controls along 5-day training. Two-way repeated-measures ANOVA with Bonferroni post-hoc correction (*P < 0.05). In (E-H), n = 3 axon-GCaMP-expressing mice; in (G, H), n = 213 avoidance trials and n = 87 failure trials; in (K, L), n = 4 control mice and 8 eNpHR-expressing mice; in (O, P), n = 8 control mice and 9 ChR2-expressing mice. In (G-J, K, L, O, P), shown as data was shown as mean ± SEM; in (F, H), shown as median. For detailed statistics information, see Supplementary Table 1.

In addition to the PVT, the PSTN also heavily projects the PBN and CeA ^22^ (Figure S6). These brain areas have been implicated in aversive learning^22,26,46^. To further characterize functional roles of the PSTN→PBN circuit and the PSTN→CeA circuit in mediating active avoidance learning, we first optogenetically activated these circuits during the CS of 2AA task (Figure S7A and F). We observed no significant differences in mean avoidance rates and mean latencies to cross the other side across 5-day training between ChR2 groups and control groups (Figure S7A-E). Furthermore, we found that optogenetic inhibition of these circuits remarkably attenuates animals’ behavioral performance during 2AA task (Figure S7F-J). These results demonstrate that the PSTN→PBN circuit and the PSTN→CeA circuits are required for active avoidance learning.

## Discussion

Although a growing body of literature has largely expanded our knowledge of neural mechanisms underlying active avoidance learning^3,5,9,14–16,42^, the hypothalamus receives less attention in the regulation of this behavior. The PSTN, a small nucleus located on the lateral edge of the posterior hypothalamus, has just been identified less than two decades and is still a relatively understudied brain region^22,23^. Despite the current research on the functional roles for the PSTN in the regulation of feeding behaviors ^24–27^, appetite suppression^28,29^, alcohol consumption^30–33^, as well as encoding of aversive experiences^26,28,35,36^, it has not been directly studied its role in the encoding and control of active avoidance learning. In the present study, through a series of well-designed experiments using animal models, we for the first time established the PSTN as a crucial brain node that encodes and controls active avoidance learning differentially via its downstream projections. This work further addresses the long-standing question of what neural substrates underlie learning within the mammalian brain.

Firstly, we identify the hypothalamic PSTN^Tac1^ neurons as novel neural substrates important for active avoidance learning. Despite recent studies indicating that the PSTN is also associated with aversion^22,37,38^, it still remains elusive about its involvement in active avoidance learning. Here, our ablation and optogenetic silencing experiments clearly demonstrated that PSTN^Tac1^ neurons are required for active avoidance learning (Figures 2 and S3), providing a first causal link these cells in the PSTN to active avoidance learning. In addition, we find that optogenetic activation of these cells is capable of promoting active avoidance learning (Figures 3 and S6). These results jointly highlight the necessity and sufficiency of Tac1-expressing cells in the PSTN in controlling active avoidance learning, adding the PSTN a new member of neural networks that control this learning^9,10,15^. Interestingly, recent studies have also demonstrated that the PSTN is involved in fear-related avoidance behaviors^26,35^. Together, these findings indicate that the PSTN involves both the regulation of avoidance behavior and its learning. Furthermore, our calcium imaging results demonstrated that punishment-predictive cues significantly enhance PSTN^Tac1^ neurons’ activity during the late phase of training (Figures 1 and S2), similar with previous findings from other brain regions, such as the lateral habenula^10^ and the PVT^9^. This result further supports the PSTN as a cellular substrate necessary for active avoidance learning.

Secondly, we find that PSTN^Tac1^ neurons could promote the generation of aversive memory. As known, learning is driven by either appetitive or negative motivational drives^4^. Both traditional tracer-based and viral-based whole-brain circuit mapping experiments revealed that the PSTN heavily connects to brain structures that involves the regulation of emotions and motivation ^22,23,27,29,34^. In our study, activation of PSTN^Tac1^ neurons can induce a strong aversive emotional drive, similar with the findings of previous studies on activation of other subpopulations of this brain region^26,28^. Remarkably, we further found that activation of these neurons can promote the generation of aversive memory (Figure 3N). This line of data further supports the role of Tac1-expressing neurons in the PSTN in active avoidance learning. On the contrary, a recent study by Serra et al. has shown that optogenetic activation of these cells in the PSTN does not promote conditioned place aversion while activation of Pitx2-expressing neurons in the subthalamic nucleus (STN), adjacent to the PSTN, mediates aversive learning^40^. Based on our calcium imaging and functional experiments (Figures 1-3), our study supports that PSTN^Tac1^ neurons play important roles in learning and memory, at least in active avoidance learning. This divergence is possibly due to the use of different conditioning parameters. Interestingly, both studies have found that PSTN^Tac1^ neurons can induce a strong aversive emotional drive^40^, strongly suggesting that the PSTN belongs to an important brain node within neuronal circuits of motivation. Moreover, the findings by Serra et al. ^40^ and our study show that both PSTN and its neighboring STN are newly identified members of neural networks controlling aversive learning, suggesting that these two brain areas function similarly in motivation and learning. In addition to *Tac1*, the PSTN also contains several other molecularly distinct cell groups (i.e., *Adcyap1* and *Crh*). Thus, examining how/whether other subtypes of the PSTN are involved in active avoidance learning represents an important future research direction of this field.

Finally, we reveal differential regulation of active avoidance learning by downstream projections of the PSTN. Despite some recent studies indicating that the PSTN projects to multiple brain regions to regulate diverse behaviors, such as feeding related behaviors^29^ and arousal^34^, it has not been directly examined the functional roles and synaptic dynamics of its downstream circuitries. Similar with previous findings ^9,10^, we observed that activity of axonal terminals of the PSTN in the PVT/IMD is gradually increased over learning. This observation resembles the expression of an LTP-like process at the PSTN-to-PVT/IMD synapses ^10^ to facilitate avoidance behavior. In addition, our functional manipulation results demonstrated that the PSTN-to-PVT circuit is both necessary and sufficient for active avoidance learning. This greatly expands our knowledge of PVT neurocircuitry in this learning^9,42^. Fascinatingly, a previous work has also identified a lateral hypothalamus-to-lateral habenula pathway involved in this learning^10^. These findings suggest the existence of two parallel neural circuits within the hypothalamus-to-thalamus circuitries for the regulation of active avoidance learning. Surprisingly, unlike the PSTN-to-PVT/IMD circuit, optogenetic activation of the PSTN→PBN circuit and the PSTN→CeA circuits did not promote active avoidance learning. Overall, these findings demonstrate that the PSTN mediates active avoidance learning possibly via different downstream projections, which is distinct from the regulation of appetite suppression by these circuits in a similar way^29^. In addition to downstream projections selected in this study, the PSTN also heavily projects to several other brain areas associated with motivational or motor components of aversive learning^22,23,29^, such as the PAG, substantia innominate (SI), and bed nucleus of the stria terminalis (BNST). Therefore, an in-depth functional study of these uncharacterized downstream circuits of the PSTN is needed.

In summary, we identified the PSTN as a key brain node for encoding and control of active avoidance learning. Importantly, instrumental defensive behaviors like active avoidance can be modified through experience and learning, resulting in that the neural circuits underlying these behaviors exhibit remarkable plasticity. Therefore, more studies are needed to fully understand the extent of the involvement of PSTN and its circuitry in active avoidance learning and neuropsychiatric disorders, as well as its potential as a therapeutic target^6^.

## Materials and Methods

### Mice

Tac1-IRES-Cre mice (8-12 weeks old) were purchased from Shanghai Model Organisms Center (Catalog # NM-KI-200083). Tac1-IRES-Cre mice were crossed to C57BL/6J mice (purchased from GemPharmatech) to produce heterozygous animals (Tac1-IRES-Cre/+, 8-16 weeks old and both sexes) for stereotaxic surgery and behavioral experiments. Animals were grouped housed in 12 h light-dark cycle (9 p.m. - 9 a.m. light), with food and water available ad libitum. Animal care and use were strictly followed according to the institutional guidelines and governmental regulations of China. Experiments were performed exactly as approved by the IACUC at Fudan University.

### Viruses

AAV9-EF1α-DIO-EGFP (Catalog #BC211231-0015-9), AAV9-CAG-FLEX-jGCaMP7f (Catalog #BC210625-0207-9), AAV9-EF1a-DIO-Axon-GCaMP6s (Catalog #BC210909-0197-9) and AAV9-EF1α-DIO-taCasp3-T2A-TEVP (Catalog #BC210909-0130-9) were purchased from Brain Case. AAV2/9-hSyn-DIO-stGtACR2-mCherry-WPRE-pA (Catalog #S0981-9-H50), AAV2/2-hEF1α-DIO-hChR2(H134R)-mCherry-WPRE-pA (Catalog #S0170-2-H50), AAV2/9-hEF1α-DIO-mCherry-WPRE-pA (Catalog #S0197-9-H50), AAV2/9-hEF1α-DIO-eNpHR3.0-mGFP-WPRE-pA (Catalog #S0178-9-H50) and AAV2/9-hSyn-DIO-EYFP-WPRE-pA (Catalog #S0276-9-H50) were purchased from Taitool Bioscience.

### Stereotaxic surgeries

Tac1-IRES-Cre/+, were anesthetized with isoflurane and mounted on a stereotaxic device (RWD Life Science). Both sexes of mice were used in the experiments. Injections were carried out using a pulled, fine glass capillary (RWD Life Science).

For fiber photometry calcium imaging, Tac1-IRES-Cre/+mice were injected with 50-100 nL AAV9-CAG-FLEX-jGCaMP7f into the PSTN (ML ±1.15 mm, AP -2.40 ∼ -2.45 mm, DV - 4.90 mm from bregma). An optic fiber (200 um core diameter, Inper) was then placed in the virus injection site in the PSTN.

For Caspase-mediated ablation of PSTN Tac1^+^ neurons, Tac1-IRES-Cre mice were injected bilaterally with 100 nL AAV9-EF1α-DIO-taCasp3-T2A-TEVP and 50 nL AAV9-EF1α-DIO-EGFP into the PSTN (ML ±1.15 mm, AP -2.40 ∼ -2.45 mm, DV - 4.90 mm from bregma).

For optogenetic activation or inhibition of PSTN Tac1^+^ neuron, Tac1-IRES-Cre mice were injected bilaterally with 100 nL AAV2/9-hSyn-DIO-stGtACR2-mCherry-WPRE-pA into the PSTN for inhibition or 100 nL AAV2/2-hEF1α-DIO-hChR2(H134R)-mCherry-WPRE-pA into the PSTN for activation (PSTN: ML ±1.15 mm, AP -2.40 ∼ -2.45 mm, DV - 4.90 mm from bregma). An optic fiber (200 um core diameter, Inper) was then placed 0.4-0.5 mm above the virus injection site in the PSTN.

For optogenetic activation of Tac1^+^ neuron in two PSTN adjacent areas (defined as P1 and P2), Tac1-IRES-Cre mice were injected bilaterally with 50 nL AAV2/2-hEF1α-DIO-hChR2(H134R)-mCherry-WPRE-pA into the P1 or P2 for activation (P1: ML ±0.75 mm, AP -2.40 ∼ -2.45 mm, DV - 4.00 mm from bregma; P2: ML ±1.70 mm, AP -2.40 ∼ -2.45 mm, DV – 5.60 mm from bregma). An optic fiber (200 um core diameter, Inper) was then placed 0.4-0.5 mm above the virus injection site in the P1 or P2.

For fiber photometry calcium imaging of the axonal terminals in the PVT from the PSTN, Tac1-IRES-Cre/+mice were injected with 100 nL AAV9-EF1a-DIO-Axon-GCaMP6s into the PSTN (ML ±1.15 mm, AP -2.40 ∼ -2.45 mm, DV - 4.90 mm from bregma). An optic fiber (200 um core diameter, Inper) was then placed in the PVT (ML 0 mm, AP -1.07 ∼ -1.43 mm, DV – 3.05 mm from bregma).

For optogenetic activation or inhibition of the downstream pathways (PSNT-PVT, PSNT-PBN, and PSNT-CeA), Tac1-IRES-Cre mice were injected bilaterally with 100 nL AAV2/9-hEF1α-DIO-eNpHR3.0-EYFP-WPRE-pA into the PSTN for inhibition or 100 nL AAV2/2-hEF1α-DIO-hChR2(H134R)-mCherry-WPRE-pA into the PSTN for activation (PSTN: ML ±1.15 mm, AP -2.40 ∼ - 2.45 mm, DV - 4.90 mm from bregma). An optic fiber (200 um core diameter, Inper) was then placed in the PVT (ML 0 mm, AP -1.50 ∼ -1.60 mm, DV - 2.95 mm from bregma) or in the PBN (ML ±1.25 mm, AP -5.50 mm, DV -3.20 mm from bregma) or in the CeA (ML ± 2.95 mm, AP - 1.40 mm, DV - 4.20 mm from bregma).

For control experiments, the animals with the same genetic background were injected with EGFP-expressing or mCherry-expressing AAVs.

### Behavioral assays

#### Automated system for quantitative analysis of two-chamber active avoidance task

In this study, we adopted and modified a widely used two-way active avoidance learning task (2AA; Fig. 1c), based on previous work^9^. The apparatus (L x W x H: 30 cm x 15 cm x 50 cm) was divided into two compartments (Fig. 1c). Electric foot shock (0.8 mA) was delivered by an Arduino-controlled Shocker (Beijing Xeye S&T Co., Ltd). Automated system for 2AA task was built on Matlab program (MathWorks). A top view of camera was used to obtain mouse real-time locations. During 2AA task, a test mouse experienced a tone (Conditioned stimulus, CS) and must shuttle to the other side of chamber in a given response window to prevent an aversive foot shock (Unconditioned stimulus, US). Each animal had a training of 100 trials each day for all 2AA task throughout the study. Each trial has a constant 10-second-long time window and the trial-trial interval is ∼ 15 s – 30 s. Once a trial starts, a sound cue is generated. If the test mouse crossed the other side of chamber within 5 seconds (defined as an avoidance trial), the sound cue was terminated immediately and no foot-shock was delivered. If the test mouse crossed the other side of chamber within 5 s – 10 s relative to trial onset (defined as an escape trial), the sound cue was terminated immediately and foot-shock was delivered 5 s after trial onset and terminated immediately when the test mouse successfully crossed compartments. If the test mouse failed to cross the chamber throughout the trial (defined as a failure trial), it experienced a maximum of shock duration (5 s) and the cue sound lasts 10 s. We examined the performance of active avoidance learning by computing the percentage of cue-guided successful shuttles over total trials and the duration from trial onset to crossing compartments (defined as the latency).

#### Caspase-3 ablation experiments

For examination of behavioral effect by caspase-3-mediated ablation before training (**Figure 2D-I**), the performance of active avoidance learning was measured described above. Caspase-3-expressing and EGFP-expressing mice were trained in 2AA task ∼3.5 weeks after viral injection. For examination of behavioral effect by caspase-3-mediated ablation after training (**Figure S3**), we first screened a cohort of Tac1-IRES-Cre mice displaying robust avoidance performance (with an average avoidance rate > 70%) after 5-day 2AA training. Those selected mice were then bilaterally injected with Cre-dependent caspase-3 (or EGFP as a control) into the PSTN. After viral expression, the mice were undergone a 2-day post-testing of 2AA task.

#### GtACR inhibition experiments

GtACR inhibition experiments (**Figure 2J-U**) were performed 2AA task described above and divided in two stimulation modes (during CS phase and US phase). To further dissect the exact role of activities of PSTN Tac1+ neurons during CS or US in controlling active avoidance behavior, optogenetic inhibition was performed during CS (continuous 5s stimulation and final blue laser output powers of 5-10 mW mm^-2^) or during US (continuous 5s stimulation and final output powers of 5-10 mW mm^-2^) from different group of test animals for each trial across the whole 7-day training. Prior to each experiment, a ferrule patch cord was coupled to the ferrule fiber implanted in the mouse using a zirconia split sleeve (Inper). An Arduino microcontroller board and a custom MATLAB program were used to control the initiation and termination of the test and record the behavioral data.

#### ChR2 activation experiments

##### ChR2 activation in the 2AA task

ChR2 activation experiments for males (**Figure 3**) and females (**Figure S4**) were performed 2AA task described above. To further dissect the exact role of activities of PSTN Tac1+ neurons during CS in controlling active avoidance behavior, optogenetic activation was performed during CS (time window: 0-5s) for each trial across a 5-day training. The stimulation parameters were as follows: frequency: 20 Hz, pulse duration: 20 ms, and laser power: ∼2.5 mW mm^-2^. Prior to each experiment, a ferrule patch cord was coupled to the ferrule fiber implanted in the mouse using a zirconia split sleeve (Inper). An Arduino microcontroller board and a custom MATLAB program were used to control the initiation and termination of the test.

##### Real time place preference test (RTPP)

ChR2-expressing or mCherry-expressing male mice were introduced into a two-chamber apparatus (60 cm x 30 cm x 30 cm for each chamber) and were allowed to freely move between the two chambers based on our recent paper^51^. Briefly, each test consisted of two consecutive 15-min sessions. In the first session, mice were allowed to freely explore the two compartments for 15 minutes, during which entering into one of the two chambers triggered optogenetic stimulation (ChR2: 473 nm, 20 ms, 20 Hz for soma stimulation, 2.5 mW mm^-2^, for up to 20-s duration). Exiting the stimulated chamber immediately terminated the light stimulation. During the second session, the opposite chamber was paired with light stimulation. An Arduino microcontroller board and a customized MATLAB program were used to control laser pulses, based on real-time tracking of mouse locations. We calculated and compared three parameters between ChR2 group and control group: time spent in the light-paired side, distances travelled in the light-paired side, and the number of visits to the light-paired side. In addition, to exclude the possibility of generating this aversive effect by sparse cells of PSTN neighboring areas, we conducted the RTPP for additional characterizations of behavioral effects of activation of sparsely Tac1-expressing neurons in two adjacent areas of PSTN (**Figure S5**).

###### Conditioned place aversion test (CPA)

To study the effect of optogenetic activation of PSTN Tac1+ neurons on generating a long-term aversive memory, we used a widely used conditioned place aversion (CPA) test. The CPA test includes three phases: pre-conditioning, conditioning, and post-conditioning. Briefly, ChR2-expressing or mCherry-expressing mice were introduced into a three-chamber apparatus to allow them to acclimate to the testing environment for 30 minutes on three days before starting the experiment. The apparatus typically consists of three interconnected chambers (the chamber on two sides: 30 cm x 30 cm x 30 cm; the middle chamber: 15 cm x 30 cm x 30 cm), each with distinct visual and tactile cues. The middle chamber is neutral and serves as a control, while the other two chambers are designated as conditioning chambers. During the pre-conditioning phase (on day 1), animals were initially placed in the middle chamber to freely explore all three chamber for 30 minutes and then the basal preference for each side was calculated for subsequent experiments. During the conditioning phase (on days 2-4), optogenetic stimulation with a cycle of 1 second light on (ChR2: 473 nm, 20 ms, 20 Hz for soma stimulation, 2.5 mW mm^-2^) and 4 second light off was consistently paired with one specific compartment that a given animal preferred to explore on day 1. During the post-conditioning phase (on day 5), animals were allowed to freely explore both chambers without any aversive stimulus present. We assessed the aversive response by calculating the percentage of time changes spent in the preferred side during pre-conditioning phase between day 1 and day 5.

#### Optogenetic manipulations of the downstream pathways in the 2AA task

ChR2 activation of the downstream circuit experiments (**Figure 4** for the PSTN-PVT; **Figure S7**) for the PSTN-to-PBN circuit and the PSTN-to-CeA circuit) was performed during the CS phase of 2AA task, but with the stimulation parameters (Frequency: 50 Hz, Pulse duration: 5 ms, and laser power: ∼2.5 mW mm^-2^). NpHR inhibition of the downstream circuit experiments (**Figure 4** for the PSTN-to-PVT circuit; **Figure S7** for the PSTN-to-PBN circuit and the PSTN-to-CeA circuit) were also conducted during the CS phase of 2AA task, but with the stimulation parameters (continuous 5s stimulation and final yellow laser output powers of 5-10 mW mm^-2^).

### Fiber photometry experiments and data analysis

Photometry experiments were performed as previously described ^47^. Fluorescence signals were acquired with a fiber photometry system (ThinkerTech). The LED power was adjusted at the tip of the optical fiber to 30–50 µW to minimize bleaching. Behaviors were recorded by a video camera. To monitor the response of Tac1-expressing cells in the PSTN (PSTNTac1) by direct foot shock, the test mouse received 10 times of 1s random aversive foot shock stimulus, with a 30∼90 s of trial-interval. To record neuronal responses of PSTN Tac1^+^ neurons during 2AA task, subject animals were allowed to freely behave in a shuttle-box chamber and the fluorescent signals were recorded simultaneously. Photometry data were analyzed using custom MATLAB programs. To measure dynamics of fluorescence intensity immediately before and after the onset of specific stimuli (e.g., sound and shock), (F−F0)/F0 (ΔF/F) was calculated, where F0 was the baseline fluorescence signal averaged over a 1 s window between 5 s and 4 s (**Figures 1**, **4E**-**H**, and **S1** and **2**), prior to stimulus onset. The analysis time window was from 2 s before CS to 10 s after CS. ΔF/F values were presented as mean with an SEM envelope. To measure neural activity (i.e., fluorescence changes of GCaMP7f or axon-GCaMP6s) of PSTN Tac1^+^ neurons or axon terminals in the PVT between early trials and late trials (pooled first 20 trials or last 20 trials of each animal on day 1) or between avoidance trials and failure trials (on day 7), we calculated the AUC per second between 0 s and 5 s from the CS onset.

### Histology and Imaging

Animals used in the study were sacrificed 4-6 weeks post-injection and perfused with 4% PFA. The brains were dissected out and fixed in 4% PFA for two additional hours at room temperature, rinsed with 3x PBS, and then put into 30% sucrose solution overnight at 4°C. To visualize viral expression and fiber placement, 60 µm sections were cut on a cryostat (RWD, China). For improving the fluorescent signals of PSTN terminals in the PVT, we conducted immunostaining experiments, largely based on our previous study^48^. Briefly, free-floating sections containing PSTN terminals in the PVT were washed in PBS 3x 5 minutes, followed by 0.5 hour blocking in 10% normal donkey serum (NDS). Primary antibodies (rabbit anti-GFP for axon-GCaMP sections or rabbit anti-mCherry for ChR2 sections; Abcam, 1:1000) were diluted in 0.5% PBS/Triton X-100 and 10% NDS and incubated overnight at 4℃. Sections were then washed with PBS 3x 5 minutes and incubated with secondary antibodies (goad anti-rabbit 488 or 555; Abcam, 1:1000) 2 hours at room temperature. Sections were then washed with PBS three times and mounted onto Superfrost Plus slides, dried > 40 minutes at room temperature, and coverslipped in 50% glycerol containing DAPI (1 ug/mL). Images were acquired using a confocal microscope (OLYMPUS FV3000) with 10x or 20x objective lens. Quantification of ablated cells (**Figures 2B** and **S3B**) and measurements of ChR2 and eNpHR special expression patterns were performed by using ImageJ software. For optogenetic experiments, we only selected data from those mice, with >90% of the infected areas within the PSTN and the optic fiber being exactly hit in the PSTN.

### Data analysis and statistics

No statistical methods were applied to pre-determine sample sizes, but our sample sizes were selected based on previous experience from related research and literature. Animals were randomly assigned to control and manipulation groups. Data collection and analysis were not performed blind to the conditions of experiments. Figures were plotted using Prism version 10 (GraphPad) or MATLAB 2023a (MathWorks). Data is reported as violin plots, or mean ± SEM plots. Statistical methods used in this study include two-way repeated measures ANOVA, one-way repeated measures ANOVA, Kruskal-Wallis test, two-sided Mann-Whitney test, two-sided unpaired t test, and two-sided paired t test (see Supplementary Table 1). The data met the assumptions of the statistical tests used. Normality of the data was tested using the Kolmogorov-Smirnov test. Statistically significant differences were established at *P < 0.05, **P < 0.01 and ***P < 0.001; ns indicates not significant. All statistic tests are summarized in the Supplementary Table 1.

## Acknowledgments

The authors are supported by the National Science and Technology Innovation 2030 Major Projects of China (STI2030-Major Projects-2022ZD0207300), National Natural Science Foundation of China (No: 82301395), and Fudan University (No: JIF2641002). We also thank Dr. Weizhe Hong from UCLA for insightful advice at initial stage of this study.

## Author contributions

Conceptualization, R.K.H. and X.H.; formal analysis, R.K.H. and R.H.; investigation, R.H., N.W., T.L., L.Z., S.L.; writing – review & editing, R.K.H. and R.H.; supervision, R.K.H.; funding acquisition, R.K.H.

## Declaration of Interests

The authors declare no competing interests.

## Data and code availability

A subset of the data and the codes used or generated in this study are available at a Zenodo repository (https://zenodo.org/records/12747284). Or directly request data and codes by contacting the corresponding author.

## Notes

### Competing Interest Statement

The authors have declared no competing interest.

